# Guessing reveals internal models of perceptual precision

**DOI:** 10.64898/2026.01.15.699528

**Authors:** Caroline Myers, Chaz Firestone, Justin Halberda

## Abstract

When observers lack sufficient information to support a confident response, they often guess. Guessing plays a pervasive role in visual cognition and working memory, yet the mechanisms that govern how observers generate guesses remain poorly understood. Standard models traditionally assume that responses produced in the absence of information are either uniformly distributed over feature space or are perhaps weighted towards prevailing environmental statistics. In contrast, here we consider an intriguing alternative: that guesses incorporate observers’ knowledge of their own perceptual capacities. We empirically measured guessing by eliciting responses under extreme target uncertainty (Experiment 1) as well as a novel “0ms presentation” approach in which no stimulus appeared but subjects believed one had (Experiment 2). We evaluated three accounts of guesses under these conditions: unsystematic (lapse) responding, biases toward environmental statistics, and a self-representational account in which guesses reflect observers’ knowledge of their own feature-dependent precision (e.g., preferring to guess feature values they believe they would be likely to miss). Guess responses were non-uniform and systematically biased toward feature values typically encoded with the least precision (e.g., oblique orientations) — a counterintuitive bias away from high-frequency, high-fidelity feature values (e.g., cardinal orientations). This complementary relationship between guessing and perceptual fidelity held within individuals and across paradigms, and was recoverable via an empirical-guess mixture model that replaced the standard uniform assumption with empirically measured guess distributions. Our findings challenge prevailing views that guesses reflect random noise, and suggest instead that guessing behavior reflects metacognitive knowledge of internal precision. Rather than defaulting to environmental priors, observers appear to model their own sensory limitations and leverage these representations to inform decisions in the absence of evidence. These results reframe guessing as a theoretically informative behavior that expresses observers’ own beliefs about their perceptual capacities.

**Significance:** Guessing is commonly treated as random noise in models of perception and memory, assumed to reflect lapses or uninformed responses. Instead, we show that human guesses are systematically structured across feature space: observers preferentially guess values they typically encode with the least precision, revealing a consistent, strategic bias away from high-fidelity representations. By directly measuring guess behavior on stimulus-absent trials and integrating these empirical distributions into a mixture model, we find that guesses on stimulus-present trials can be systematically recovered, and that they too form the complement of perceptual precision. These findings challenge foundational psychophysical modeling assumptions and position guessing as a strategic, informative behavior that engages self-representation.

## MAIN TEXT

When we don’t know what we’ve seen, we often guess. This is familiar in daily life and ubiquitous in behavioral experiments that probe perception and memory. Despite its central role, guessing is typically treated as theoretically uninformative. Standard models assume that responses produced in the absence of perceptual information are random, approximated by a uniform distribution over feature space and captured by a lapse parameter. This assumption underlies widely used mixture models and shapes how guess rates are interpreted across behavioral, neural, and computational studies (Prins 2012; Klein, 2001; Zchaluk & Foster, 2009; Wichmann & Hill, 2001), but it has never been tested directly.

In principle, guessing need not be random. At least three distinct computations are possible. Guess responses could be unsystematic, reflecting arbitrary responding when information is absent. Alternatively, guessing could be systematic in a manner consistent with environmental priors, such that observers leverage knowledge that some feature values are more probable in the world than others (e.g., a preference for cardinal orientations, which are more prevalent in natural scenes; Girshick, Landy, & Simoncelli 2011; Harrison, Bays & Rideaux, 2023; Oliva 2001; Hansen & Essock 2004; Girshick, Simoncelli & Landy, 2011; Appelle, 1972). A third possibility, central here, is that guessing reflects self-representation. On this view, observers possess internal knowledge of how precisely different feature values are internally represented, and strategically condition their guesses on this knowledge when perceptual evidence is insufficient. Guessing would therefore be structured by feature-dependent precision, rather than by external stimulus statistics.

This self-representational account makes a distinctive and counterintuitive prediction. When forced to guess, observers should avoid feature values they normally represent with high precision and instead preferentially sample from low-precision regions of the feature space. In orientation, this predicts an increased tendency to guess oblique orientations and a systematic avoidance of cardinal orientations, despite the fact that cardinals are both perceptually advantaged and more prevalent in natural scenes. Such a pattern would contradict environmental-prior accounts and instead imply that guess behavior reflects metacognitive knowledge of one’s own perceptual limitations.

Here, we tested these alternatives in orientation report tasks using two complementary approaches. In Experiment 1, we elicited guess-like responses under extreme target uncertainty using extremely high load and brief exposure trials, isolating responses that are effectively untuned to the target. In Experiment 2, we embed backward-masked 0-ms trials in which no stimulus is presented, providing an observer-specific empirical measure of pure guessing. These empirical guess distributions allow us to test whether guessing is uniform, prior-driven, or structured by internal precision. We further use these distributions to constrain a two-component mixture model, replacing the standard uniform guess term with each observer’s measured guess density. This approach yields trial-wise posterior estimates that distinguish responses likely driven by target-centered internal representations from those best characterized as guesses, enabling a direct test of whether guessing and perceptual precision exhibit a systematic relationship.To preview our results, we found that guesses were reliably non-uniform and biased toward feature values observers typically represent with the least precision. Thus, guessing behavior provides a direct readout of internal models of representational fidelity, revealing that observers not only monitor their own perceptual limitations, but actively exploit this knowledge to guide behavior when evidence is absent.

## RESULTS

### Experiment 1

In Experiment 1, (*n* = 10) observers completed a continuous-report working memory orientation task with two interleaved two trial types designed to separate high-fidelity encoding from guess-like responding: low-load, long-duration “precision” trials (1 item, 1000 ms) and high-load, ultra-brief “guess” trials (36 items, 16 ms; see **SI Methods**).

#### Precision and guess trials engage distinct response regimes

Responses on 1-item, 1000ms precision trials tightly tracked the target orientation (Pearson r = .892, 95% bootstrap CI [0.862, 0.922]., p< .0001; **Figure 2A**). By contrast, responses on 36-item, 16 ms guess trials were essentially untuned to the target (Pearson r = -.011, 95% bootstrap CI [−0.0582, 0.0336], p > .5; **Figure 2B**). The within-observer difference was reliable (mean Δr = 0.908, CI [0.863, 0.949]; exact sign test p < 0.001).

**Figure 1.**
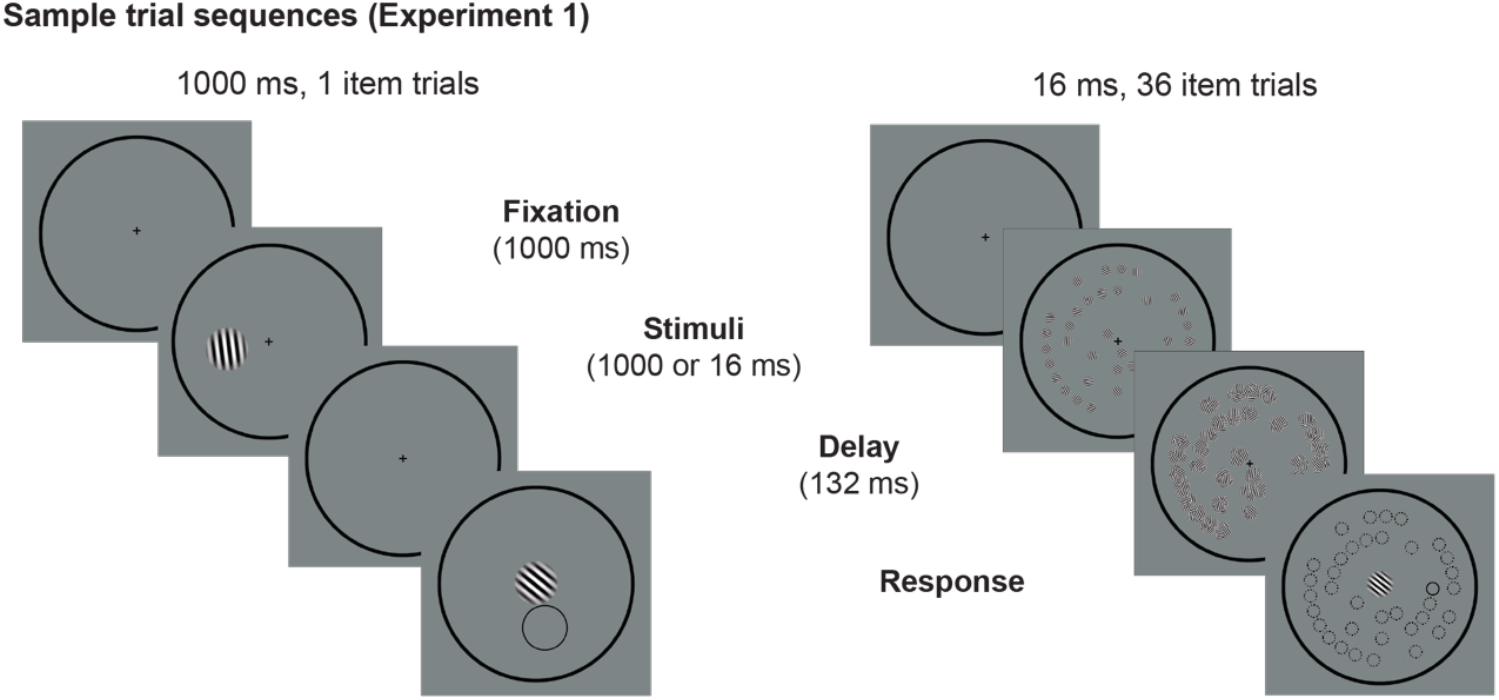
Sample trial sequences for the two trial types used in Experiment 1. Each trial began with a centrally presented fixation cross, which participants clicked to initiate the trial. A circular response ring remained visible throughout. On precision trials (left), a single oriented Gabor patch was presented for 1000 ms at one of nine possible peripheral locations. Following stimulus offset, participants adjusted the orientation of a centrally presented response Gabor to match the remembered target orientation. On guess trials (right), thirty-six Gabors (one target and 35 distractors) were presented simultaneously for 16 ms, followed by a 132-ms backward mask composed of randomly oriented Gabors. At response, the target location was indicated by a solid outline and distractor locations by dashed outlines. In both trial types, responses were made using a continuous-report procedure, and accuracy was emphasized over response speed.

**Figure 2.**
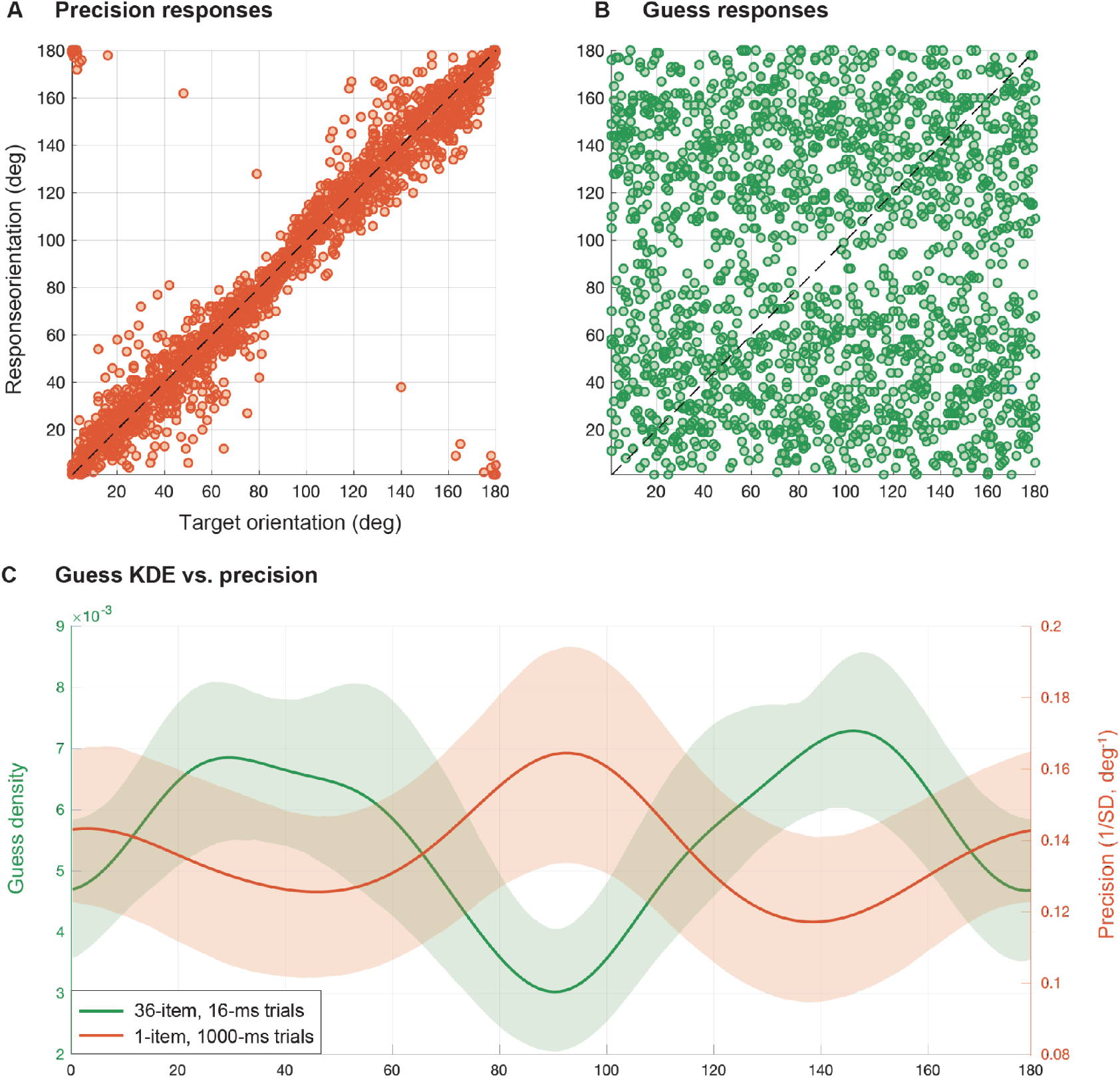
Guesses are non-uniform, forming the complement of precision. **(A)** Pooled target-response correlations on precision trials (set size 1, 1000ms), showing strong target tuning; dashed unity line indicates perfect performance. **(B)** Subject-wise target-response correlations on guess trials (set size 36, 16ms), showing near-zero tuning in the minimal-information condition. **(C)** Guess response probability (left axis) and precision landscape (right axis; precision = 1/SD of signed error, deg^−1^) overlaid as a function of absolute orientation. Guess distributions were estimated per observer using circular KDE; precision landscapes were estimated from precision trials using Gaussian weighting over target orientation (see **SI Methods**). Across observers, guess density and precision were reliably anticorrelated (mean r = −0.335857; sign-flip p(left) = 0.007805). For visualization, both curves were additionally smoothed with the same circular Gaussian kernel (σ = 10 deg), matching smoothness across curves; shaded bands show 95% bootstrap CIs across observers.

#### Guess responses are non-uniform across orientation space

To characterize guess behavior, responses from 36-item/16-ms trials were collapsed across targets (to which they were unresponsive) and fit using a circular kernel density estimate over the 180° axial orientation space. If guessing were a pure random process, the distribution of responses on guess trials should be approximately uniform. Instead, guess response distributions deviated from uniformity in 9/10 observers (Kuiper permutation p < 0.05; **Figure 2C**).

#### Guessing forms the complement of precision

We next tested whether guessing relates systematically to an observer’s own feature-dependent precision. Each observer’s precision landscape was estimated from 1-item/1000-ms trials as the inverse of the smoothed SD of signed error as a function of target orientation (**SI Methods**). If guess behavior were driven by an environmental prior favoring cardinals, the guess distribution should peak near those high-precision axes. In contrast, a self-representational account predicts that guess density should be highest where the observer’s precision is lowest– i.e., that guessing should align with the complement of the precision landscape.

Guess densities were compared to precision landscapes at their true physical alignment (relative to the cardinal axes), using an exact circular-shift alignment test that preserves smoothness while breaking physical correspondence (SI Methods). Guess density and precision were anticorrelated (mean r = −0.336, 95% CI [−0.519, −0.132]; sign-flip p(left) = 0.007805; **Figure 2C**), consistent with guesses concentrating in orientations that the same observer represents least precisely.

### Experiment 2

#### Pure guessing is structured and supports empirical-guess mixture modeling

Experiment 1 used conditions intended to induce guess-like states, but weak stimulus traces could in principle still influence responding even under extreme load and brief exposure. Experiment 2 therefore embedded backward-masked 0-ms (stimulus-absent) trials among stimulus-present trials varying set size (1, 3, 5) and display duration (16– 300 ms; **SI Methods**).

These stimulus-absent trials both (1) provide an observer-specific measure of pure guessing and (2) enable a stronger test of the complement account: if guessing reflects self-represented precision limitations, complementarity should be evident not only on stimulus-absent trials, but also in the subset of stimulus-present trials on which observers effectively guessed.

#### Stimulus-absent trials yield a non-uniform empirical guess distribution

Responses on 0-ms trials were systematically non-uniform in 10/10 observers (within-observer Kuiper permutation test p < 0.05; **Figure 3**). Moreover, each observer’s 0-ms guess density was positively similar to that same observer’s induced-guess density from Experiment 1 (mean within-observer *r* = 0.444, 95% CI [0.199, 0.635]; sign test p = 0.0107), consistent with a shared guessing strategy across paradigms.

**Figure 3.**
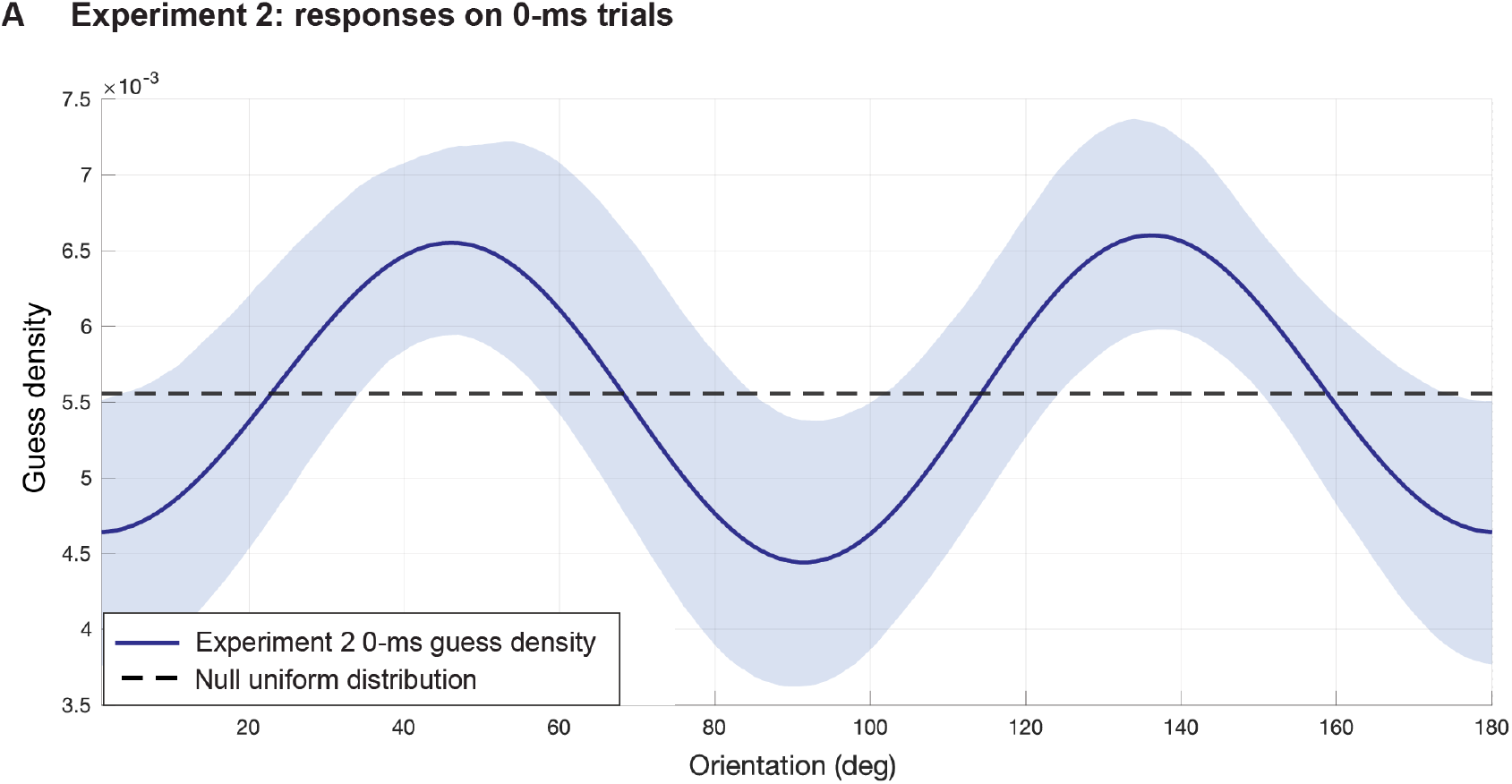
Experiment 2 0-ms trial guess response structure. Responses on 0-ms guess trials were distinctly non-uniform in all subjects. Each observer’s 0-ms guess density was positively correlated to that same observer’s induced-guess density from Experiment 1 (mean within-observer *r* = 0.444, 95% CI [0.199, 0.635]; sign test p = 0.0107), consistent with a shared guessing strategy across paradigms.

#### Empirical-guess mixture model

Standard continuous-report mixture models treat guesses as uniformly distributed over the response space, and attempt to control for guessing via the inclusion of a parametric lapse parameter. The preceding results show that this assumption is violated in a reliable, structured way. More importantly, they raise a deeper inferential problem: when guessing is structured in absolute feature space, collapsing responses into target-centered error distributions obscures this structure and renders guessing statistically indistinguishable from extreme imprecision. Can guess behavior be empirically identified and separated from stimulus-driven representations without discarding the structure it exhibits across feature values?

We fit a two-component mixture model to target-present trials in Experiment 2 (**Figure 4A)**, with (1) an internal component defined by a normalized target-centered distribution whose SD varies with target orientation according to a two-parameter symmetric oblique-effect function and (2) a nonparametric guess component fit to the observer’s measured 0-ms guess KDE **(SI Methods)**.

**Figure 4.**
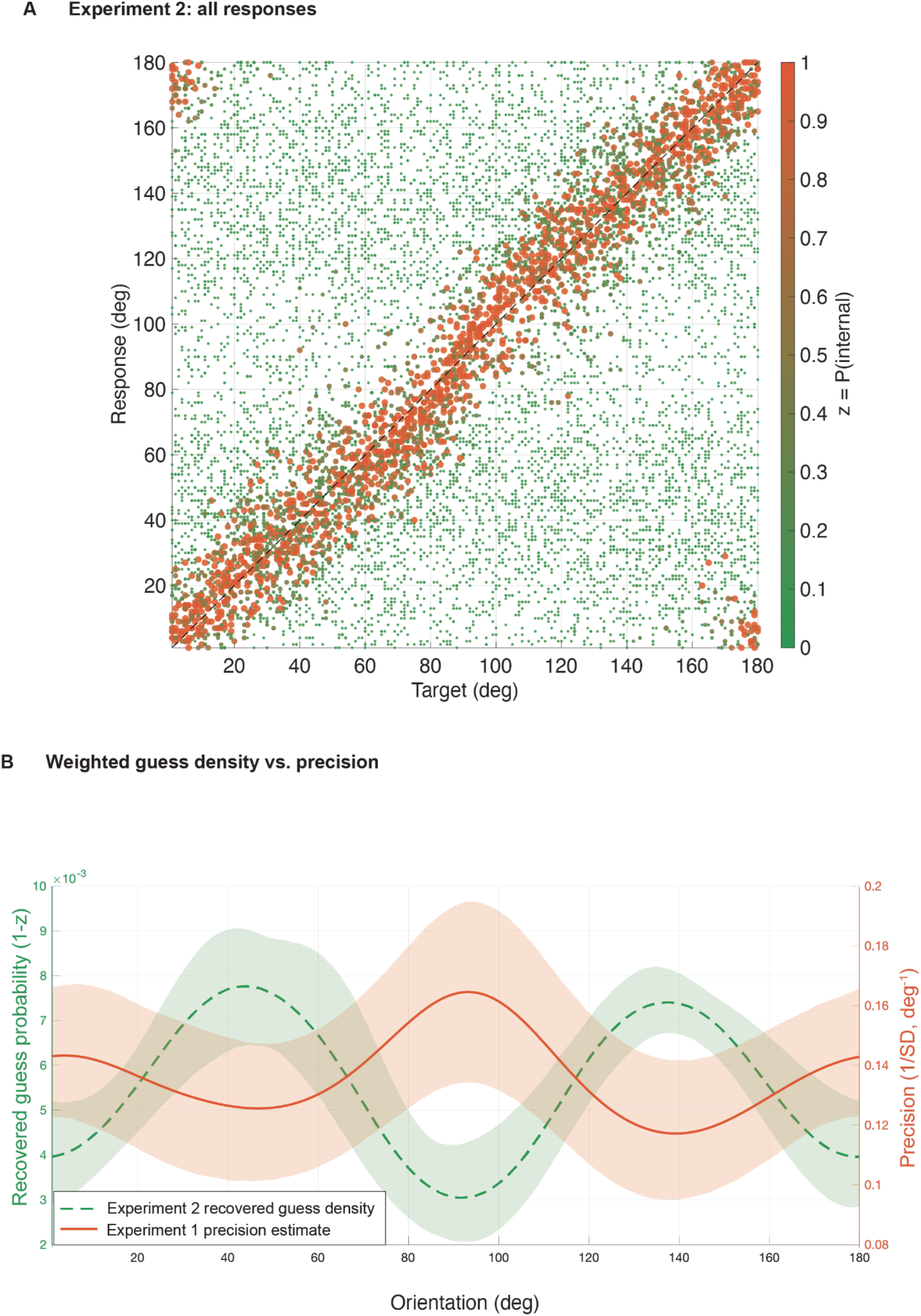
Recovered guess structure on stimulus-present trials tracks the complement of independently measured precision. (A). Experiment 2, all responses. Target orientation (x-axis) is plotted against response orientation (y-axis) for all stimulus-present trials in Experiment 2, pooled across set sizes and display durations. Each point is a single trial, colored by the empirical-guess mixture model’s trial-wise posterior probability that the response was generated by the internal (target-driven) component, *z*_*i*_ (color bar). Warmer colors indicate higher *z*_*i*_ (responses more consistent with a target-centered internal representation); cooler colors indicate lower *z*_*i*_ (responses more consistent with the empirical guess component derived from 0-ms trials). The dashed diagonal indicates perfect target–response correspondence. **(B)** Complementarity between recovered guess density and precision, measured independently on Experiment 1 1000ms, 1 item trials. Dashed curve (left y-axis) shows the group mean recovered guess density from Experiment 2 stimulus-present trials, reconstructed by weighting each response by its posterior probability of guessing, 1−z_i_ from the empirical-guess mixture model and estimating a circular KDE over the 180° axial orientation space. The solid curve (right y-axis) shows the group mean precision landscape from Experiment 1 precision trials, computed as 1/SD(θ), where SD(θ) is the smoothed target orientation-dependent standard deviation of signed error.

#### Trial-wise posteriors recover structured guessing on stimulus-present trials

To test whether the same structured guessing measured on 0-ms trials also operates when a stimulus is physically present, we reconstructed a guess-weighted response density from stimulus-present trials by weighting each response by its posterior probability of guessing and then estimating a circular KDE on these weighted responses. The recovered guess density from stimulus-present trials closely matched the independently measured 0-ms guess density (mean r = 0.869, 95% bootstrap CI [0.836, 0.902]; **Figure 4B**), indicating that the model’s trial-wise separation recovers empirical guess structure from stimulus-present data.

We also quantified the expected recovered *q*_0_ similarity under a null in which stimulus-present guesses are uniform across orientation. Specifically, for each observer we simulated stimulus-present responses preserving the fitted internal component and trial structure, but drawing guess responses from a uniform distribution; we then applied the identical posterior-weighted KDE recovery procedure. All observers’ recovered–*q*_0_ correlations exceeded this uniform-guess null (*p* < . 0.01). indicating that the recovered– *q*_0_ match is not an artifact of using *q*_0_ in the posterior.

#### Guessing on stimulus-present trials is biased toward low-precision orientations

We then tested the key theoretical prediction: if guessing reflects self-representational knowledge of perceptual limitations, guess-like responses on stimulus-present trials should be biased toward the least precise orientations and away from the most precise orientations. Using the recovered (1−*z*)-weighted guess density from stimulus-present trials, complementarity with the independently measured Experiment 1 precision landscape was reliably negative across observers (mean r = -0.495, 95% bootstrap CI [−0.669, −0.311]; one-sided sign-flip test p(left) = 0.00195). Thus, on the subset of stimulus-present trials where observers behave as if they are guessing, their responses are systematically biased toward the least precise regions of their internal feature space.

## DISCUSSION

Across two experiments, we directly measured guess behavior under conditions of extreme uncertainty (Experiment 1) and on completely stimulus-absent trials (Experiment 2) and found that guess responses were reliably non-uniform. Guess density varied systematically with orientation in a manner that closely tracked the inverse of each observer’s precision landscape. Observers preferentially guessed orientations they represented least precisely and avoided orientations they represented most precisely. This inverse relationship was evident within individual observers and was recovered independently from both stimulus-absent trials and guess-like responses on stimulus-present trials.

These findings bear directly on how continuous-report data are modeled and interpreted. In standard discrete-state working-memory mixture models, responses are decomposed into an internal-representation component and a guess component assumed to be uniform over feature space (Zhang & Luck, 2008; Rouder et al., 2008; Adam, Vogel & Awh, 2017). Variable precision models reject this dichotomy, treating all responses as noisy but meaningful, and positing a continuous distribution over representational fidelity (van den Berg et al., 2012; Schurgin, Wixted & Brady, 2020). In these models, large errors are assumed to arise from extremely low precision, rather than the absence of memory altogether.

Our results place new pressure on both classes of models. The defining assumption shared by both is that guesses are unstructured: either uniform (in the discrete case), or non-existent (in the continuous case). But when guess behavior is structured, this assumption fails in principled ways.

Our results show that when internal signal is absent, guessing is not uniform, and when signal is present but degraded, guess-like responses still follow orientation-dependent patterns that are unrelated to the stimulus shown. These patterns are consistent across stimulus-absent and stimulus-present conditions and are defined in orientation space, not error space. That is, the structure in guessing is organized around internal feature representation, not the stimulus target.

This introduces a problem of misattribution. For discrete models, the estimated guess weight no longer isolates trials without information, because responses produced under guessing are not exchangeable across feature values. As a result, orientation-dependent regularities in guessing may be misinterpreted as properties of the internal representation, such as reduced precision or increased variability. For continuous models, this poses a deeper problem: The assumption that even guess responses reflect degraded signal becomes difficult to maintain once systematic biases appear even in the absence of a stimulus.

In both cases, the presence of orientation-specific structure in guess responses introduces parameter ambiguity. Model fits may conflate true imprecision with feature-dependent bias in guessing, echoing prior critiques of how mixture model outputs can be distorted by misattributed structure in error distributions (Panichello, DePasquale & Buschman, 2019; Bays, 2014). This misattribution is compounded when responses in continuous-report tasks are collapsed into target-centered error distributions. This effectively removes absolute feature information and projects all responses into a space defined as relative to the target. A non-uniform distribution of responses that are truly independent from any target (e.g., guesses) will therefore appear as uniform noise in this error space, not because the responses lack structure, but because the structure is defined in absolute feature space rather than relative to the target. Models fit to such collapsed error distributions may therefore appear to support uniform guessing or extremely low-precision representations, not because those assumptions are warranted by the data, but because the preprocessing step itself renders alternative explanations impossible. In this sense, common modeling conclusions about guessing and precision reflect how the data are transformed rather than how observers actually respond. This also reflects a deeper modeling assumption: that responses without target correspondence lack meaningful structure. Our results directly challenge that assumption.

We resolve this tension by introducing an empirical-guess mixture model that replaces the uniform guess term with each observer’s measured guess distribution. This model both (1) captures non-uniform, structured guessing, without introducing new parameters, and (2) provides posterior estimates that separate stimulus-present trials based on whether they reflect an internal representation of the stimulus or informed guessing. Indeed, replacing the uniform guess term with each observer’s empirically measured 0-ms guess distribution produced a large and consistent improvement in model fit in all observers, indicating that structured guessing captures systematic variance missed by standard lapse assumptions.

This raises a natural question about the computational role of guessing: What does it mean to guess strategically? Our findings suggest that even in the absence of perceptual evidence, observers draw on internal knowledge of their own perceptual limitations to constrain responses to likely guesses. In orientation space, this manifests as a systematic bias toward oblique orientations where perceptual precision is poor.

This pattern runs counter to that predicted by most standard Bayesian observer models, which emphasize the role of environmental regularities in shaping perceptual estimates (Wei & Stocker, 2015; Stocker & Simoncelli, 2006). For instance, observers are typically biased toward cardinal orientations, a pattern long attributed to their statistical prevalence in visual environments (Girshick et al., 2011; Harrison, Bays & Rideaux, 2023; Coppola et al., 1998; Howe & Purves, 2005; Hansen & Essock, 2004). These environmental anisotropies are mirrored in perceptual performance, where thresholds for detecting or discriminating cardinal orientations are consistently lower than for oblique ones, a well-established phenomenon known as the oblique effect (Appelle, 1972; Essock, 1980; Heeley & Timney, 1988; Westheimer 2003).

Importantly, these behavioral asymmetries also align with anisotropies in neural tuning: cardinal preferences are reliably observed in early visual cortex, including in coarse-scale population coding in V1 (Roth, Kay & Merriam, 2022) and orientation-selective BOLD responses (Patten, Mannion & Clifford, 2017), supporting a model in which neural architecture and perceptual priors are jointly shaped by environmental regularities (Geisler, 2008).

However, our results show that when external structure is absent or unavailable, observers default not to what is likely in the world, but to what is likely given their own representational limitations. This is not a prior over the stimulus, but a bias conditioned on introspective uncertainty that cannot be captured by models that conflate encoding priors with perceptual likelihoods (even those that unify efficient coding with Bayesian decoding; Wei & Stocker, 2015). In those models, repulsive perceptual biases arise from asymmetric likelihoods shaped by natural stimulus statistics. But our data reveal repulsive biases in the absence of a stimulus altogether, suggesting that these patterns reflect internal models of sensory fidelity rather than environmental probability.

A guessing bias conditioned on introspective uncertainty may be more consistent with theories of resource-rational inference, in which decision strategies are shaped by internal constraints as much as by external statistics (Lieder & Griffiths, 2017; Prat-Carrabin et al., 2024; Van den Berg & Ma, 2018). Rather than optimize guess responses for environmental likelihood (e.g., guessing high-prevalence cardinal orientations), observers optimize for representational plausibility: when uncertain, they report what they expect themselves to miss. This form of introspective strategy is not captured by standard Bayesian accounts, which only assume access to environmental priors, nor by more general models that deny metacognitive access to internal noise.

Instead, it reflects a form of pragmatic metacognition, where responses are generated based on knowledge of one’s own uncertainty. This interpretation is supported by emerging work suggesting that confidence and precision can be tracked even in the absence of external stimuli, and that metacognitive judgments may rely on internal monitoring of representational fidelity (Fleming & Daw, 2017; Boldt, de Gardelle & Yeung, 2017). Our results extend these findings by demonstrating that such monitoring influences primary task responses.

This complementarity links guessing to self-knowledge about perceptual representation. When perceptual evidence was insufficient to support an internal representation of the target, responses were biased toward orientations associated with lower precision. This supplies a behavioral route to metacognition in a domain where measurement is typically indirect. Standard approaches rely on confidence judgments or other secondary reports that are vulnerable to both response bias and the tight coupling between performance and awareness, making it difficult to isolate metacognitive knowledge from first-order task demands (Lau, 2008; Morales, Chiang, & Lau, 2015). Moreover, existing measures necessarily depend on the presentation of a stimulus, introducing additional variables that complicate interpretation (Morales, Odegaard, & Maniscalco, 2022). In our tasks, observers’ beliefs about their own precision are expressed in the primary report itself. The structure of guessing therefore provides a quantitative readout of observers’ models of their own perceptual limitations, without requiring explicit report or relying on stimulus-driven responses.

In sum, guessing is structured, feature-specific, and systematically related to perceptual precision. Measuring that structure reveals a complementary relationship between guessing and precision and provides a new tool for metacognitive inference.

More broadly, our results challenge the assumption that guess responses reflect failure. When observers lack evidence, they still respond in ways that reflect internal models of error and representational uncertainty. Recognizing this reframes guessing as a theoretically informative behavior and establishes a new approach for probing internal models of perceptual representation without relying on confidence judgments or explicit report.

