## Supplementary material for "Guessing reveals internal models of perceptual precision": SI_Methods

### SUPPLEMENTARY MATERIALS AND METHODS

#### Participants

Ten participants participated in both experiments in a fully within-subjects design. Participants were recruited via campus recruitment and the Johns Hopkins University SONA subject pool and received course credit for completion of one-hour in-house sessions. All participants reported normal or corrected-to-normal vision. Study procedures were approved by the Johns Hopkins University Institutional Review Board.

#### Apparatus and software

All stimuli were generated and presented using custom JavaScript in conjunction with the jsPsych framework (de Leeuw, 2015) and run locally on a high–refresh-rate LCD monitor (nominal refresh rate: 120 Hz) in a dark, sound-attenuated testing room. Viewing distance was fixed at 57 cm using a chinrest. The display was calibrated and luminance-linearized using a vPixx i1Display Pro Spectra-Colorimeter photometer. All preprocessing, statistical analyses, and modeling were conducted using custom-written MATLAB code (R2017b).

#### Stimuli

Stimuli in both experiments were sinusoidal Gabor patches presented on a uniform gray background. Patches were windowed by a circular Gaussian envelope and rendered at full Michelson contrast. Orientation was defined over a 180° axial space, such that orientations separated by 180° were treated as equivalent. On trials with multiple items, orientations were sampled without replacement subject to a minimum angular separation of 5° to avoid identical or near-identical orientations.

#### Experiment 1: Guess-like responding under extreme uncertainty

#### Task overview

Participants completed a continuous-report orientation memory task designed to contrast high-fidelity encoding with guess-like responding under extreme uncertainty. Each participant completed a single one-hour session comprising 360 total trials. Two trial types were randomly interleaved throughout the session: precision trials (50%) and guess trials (50%).

#### Trial sequence

Each trial began with a centrally presented fixation cross. Participants initiated the trial by clicking the fixation cross with the mouse. A black circular response ring remained visible throughout the trial. Responses were made by rotating the orientation of a centrally presented Gabor patch using the mouse and confirming the response with a mouse click. Accuracy was emphasized over response time.

**Precision trials**
On precision trials, a single oriented Gabor patch (180 px diameter) was presented for 1000 ms at one of nine equally spaced locations on an imaginary circle centered on fixation (radius: 320 px). Target orientation was sampled uniformly from the 180° orientation space. Following stimulus offset, the response display appeared immediately. The response display consisted of an adjustable central Gabor patch and a solid black outline marking the target location. Participants adjusted the central Gabor to match the remembered target orientation and confirmed their response with a mouse click. The starting orientation of the response Gabor was randomized on each trial to minimize motor and response biases. Accuracy was emphasized over speed.

**Guess trials**
Guess trials were designed to elicit responses under conditions of extreme perceptual uncertainty while preserving the general structure of precision trials. On each guess trial, thirty-six Gabor patches (one target and 35 distractors; each 45px diameter) were presented simultaneously for 16 ms. Stimuli were positioned at locations sampled without replacement from a fixed set of 91 possible spatial positions distributed across four concentric rings centered on fixation (radii: 120, 200, 280, and 360px). Immediately following stimulus presentation, all items were masked for 132 ms by spatially overlapping, randomly oriented Gabor patches drawn uniformly from the stimulus space (90 X 90px). At response, the target location was indicated by a solid black outline, and distractor locations were indicated by dashed outlines to minimize spatial uncertainty. Participants reported the target orientation using the same continuous-report procedure as on precision trials.

**Experiment 2: Stimulus-absent guessing and empirical-guess modeling**

**Task overview**
Experiment 2 extended the task used in Experiment 1 to directly measure guessing and to support model-based separation of guessing from target-driven responses. Participants completed four one-hour in-house sessions comprising 3600 total trials. Apparatus, response method, and general task structure were identical to Experiment 1.

**Trial sequence**
Each trial began with a centrally presented fixation cross. Participants initiated the trial by clicking the fixation cross with the mouse. A black circular response ring remained visible throughout the trial. Responses were made by adjusting the orientation of a centrally presented Gabor patch using the mouse and confirming the response with a mouse click. Accuracy was emphasized over response speed.

**Stimulus-present trials**
On stimulus-present trials, oriented Gabor patches (90px diameter) were presented at set sizes of 1, 3, or 5 items for one of four display durations (16, 66, 132, or 300 ms).

Stimuli were positioned at locations sampled without replacement from a fixed set of 24 possible spatial positions distributed across two concentric rings centered on fixation (radii: 120 and 300 px).

Following stimulus offset, all trials were immediately followed by a 132-ms backward mask composed of randomly oriented Gabors drawn uniformly from the stimulus space. Following the masks, the response display appeared. The target location was indicated by a solid outline: dashed placeholders indicated locations of distractors to minimize location uncertainty at the time of response. Participants reported the target orientation using the same continuous-report procedure as in Experiment 1.

**Stimulus-absent trials**
On stimulus-absent (0-ms) trials, no stimulus was presented prior to the mask. These trials were otherwise identical to stimulus-present trials, including masking and response displays, and were randomly interleaved with stimulus-present trials. Participants were not informed of the presence of stimulus-absent trials. Participants were probed following successful completion of the experiment: none none spontaneously identified the purpose of the study none spontaneously identified the purpose of the study, the presence of the 0-ms trials, or reported any strategies consistent with deliberate or strategic guessing.

### QUANTIFICATION AND STATISTICAL ANALYSIS

#### Experiment 1

**Axial orientation transformation**
All analyses were conducted in an axial 180° orientation space (i.e., θ and θ+180° are equivalent). We represent orientations on a 1° grid:

$$r \in \{1,2,\dots,180\},$$

Continuous (non-integer) target and response angles are wrapped to this axial domain by:

$$\mathrm{wrap}_{180}(x) \;=\; ((x-1)\,mod\,180)+1,$$

thus mapping values into [1, 180] while preserving degrees.

The signed axial difference (response error) between response $r$ and target $t$ was defined as:

$$\Delta(r,t) \;=\; ((r-t+90)\,mod\,180)-90, \Delta(r,t) \in [-90°, 90°).$$

We also use the corresponding axial distance for computations in which only error magnitude matters as:

$$\delta(r,t) = |\ \Delta(r,t)\ | \in [0°, 90°],$$

**Circular KDE for guess densities**

Circular guess densities $q(r)$ were estimated over the 180° axial orientation space using a circular Gaussian kernel density estimate (KDE) evaluated on the 1° grid $r \in \{1, \ldots, 180\}$. Given a set of orientation responses $\{r_j\}_{j=1}^{N}$, we computed the unnormalized guess density as:

$$\tilde{q}(r) = \sum_{j=1}^{N} \exp\left[-\frac{1}{2}\left(\frac{\delta(r, r_j)}{h}\right)^2\right],$$

with bandwidth $h = 3°$. The KDE is then normalized to unit mass on a grid:

$$q(r) = \frac{\tilde{q}(r)}{\sum_{r'=1}^{180} \tilde{q}(r')}, \sum_{r=1}^{180} q(r) = 1.$$

**Non-uniformity**

Within each observer, deviation from uniformity was tested with Kuiper's statistic on the empirical CDF of responses mapped to [0, 1). Significance values were computed by permutation against $n_{perm}$ = 20,000 samples of matched size.

**Precision landscape**

Precision landscapes were estimated from precision trials as the inverse of the target-dependent standard deviation of signed axial error. For precision trial $i$, signed error was defined as:

$$e_i = \Delta(r_i, t_i) \in [-90°, 90°),$$

where $t_i$ was the target orientation and $r_i$ is the response orientation.

For each orientation bin $\theta \in \{1, \ldots, 180\}$, we computed Gaussian weights over target orientation using axial distance:

$$d_i(\theta) = \min\left(|\ t_i - \theta\ |,\ 180 - |\ t_i - \theta\ |\right), \qquad w_i(\theta) = \exp\left[-\frac{1}{2}\left(\frac{d_i(\theta)}{\sigma}\right)^2\right],$$

with $\sigma = 10°$. The weighted mean and SD of error are:

$$\mu(\theta) = \frac{\sum_i w_i(\theta) e_i}{\sum_i w_i(\theta)}, SD(\theta) = \sqrt{\frac{\sum_i w_i(\theta)\,(e_i - \mu(\theta))^2}{\sum_i w_i(\theta)}}$$

Precision is defined as:

$$P(\theta) = \frac{1}{SD(\theta)} (\mathrm{deg}^{-1}).$$

**Complementarity and exact circular-shift tests**
Complementarity was assessed within each observer as the Pearson correlation between the observer's guess density and precision landscape at their true physical alignment in absolute orientation space.

Let $q(r)$ denote the observer's guess KDE and $P(\theta)$ denote the observer's precision landscape (both defined on the same 1° grid). To match smoothness across the two curves for correlation and alignment testing, both curves were additionally smoothed with the same circular Gaussian kernel ($\sigma = 10°$; truncated at $\pm 4\sigma$ and normalized), producing $q_{\mathrm{smooth}}$ and $P_{\mathrm{smooth}}$.

The observed complementarity statistic is the correlation at true alignment:

$$r_{\mathrm{obs}} = \mathrm{corr}(q_{\mathrm{smooth}}, P_{\mathrm{smooth}}).$$

To test whether this correlation depended on physical alignment (rather than arbitrary phase matching induced by smoothing, which renders nearby neighboring points non-independent), we constructed an exact circular-shift null by shifting one curve by all non-zero offsets $k \in \{1, \ldots, 179\}$ and recomputing the correlation:

$$r_k = \mathrm{corr}\,(\mathrm{circshift}(q_{\mathrm{smooth}}, k), P_{\mathrm{smooth}}).$$

For a **left-tailed** complementarity test (negative correlation), the exact p-value is computed as:

$$p_{\mathrm{left}} = \frac{\#\{k: r_k \le r_{\mathrm{obs}}\} + 1}{180}.$$

Group-level confidence intervals over observers were computed by bootstrapping observers (5,000 resamples). We computed group-level p-values for a mean correlation with a one-sided sign-flip test on observer-level correlations.

**Experiment 1 target-response tuning (precision vs. guess trials)**
Within each observer, target–response tuning was quantified as the Pearson correlation between target orientation and response orientation on precision trials and on guess trials. For group-level inference on the within-observer difference in tuning $(r_{\mathrm{precision}} - r_{\mathrm{guess}})$, we used an exact sign test on the number of observers showing stronger tuning on precision trials.

**Experiment 2 and mixture model**

**Model definition**
We fit, within each observer, a two-component mixture model to stimulus-present trials in Experiment 2. For trial $i$, let $t_i$ be target orientation, $r_i$ response orientation, and $c_i$ index the condition (set size X display duration). The response likelihood is:

$$p(r_i \mid t_i, c_i) \; = \; w_{c_i}\, p_{\text{int}}(r_i \mid t_i; a, b) \; + \; (1 - w_{c_i})\, q_0(r_i),$$

where:

(1) $q_0(r)$ is the observer's fixed 0-ms KDE (unit mass on the 1° grid: $\sum_{r=1}^{180} q_0\,(r) = 1$); in likelihood computations, $q_0(r_i)$is evaluated at continuous $r_i$by circular linear interpolation on the 1° grid.
(2) $w_c \in [0,1]$ is the condition-specific weight of the internal (target-centered) component.
(3) $p_{\text{int}}$ is the internal component, defined as a target-centered axial Gaussian whose width depends on target orientation via a two-parameter "bowtie" function.

The internal component is defined as:

$$p_{\text{int}}(r_i \mid t_i; a, b) = \frac{\exp\left[-\frac{1}{2}\left(\frac{\delta(r_i, t_i)}{SD(t_i; a, b)}\right)^2\right]}{\sum_{r=1}^{180} \exp\left[-\frac{1}{2}\left(\frac{\delta(r, t_i)}{SD(t_i; a, b)}\right)^2\right]},$$

where $\delta(\cdot,\cdot)$ is axial distance and the orientation-dependent SD is:

$$SD(\theta; a, b) = \frac{a + b}{2} - \frac{a - b}{2}\cos\,(4\theta).$$

(Here, $\theta$ is in degrees; equivalently $\cos\,(4\theta)$can be written $\cos\,(4\theta\pi/180)$. This parameterization yields $SD = a$ at obliques and $SD = b$ at cardinals; during fitting we enforced $a \geq b$, consistent with known anisotropies in human precision across orientation space (e.g., Hansen & Essock 2004, Girshick, Landy & Simoncelli 2011, Appelle, 1972).

**EM fitting and posteriors**
Parameters were fit via an EM algorithm with $a, b$ shared across conditions within each observer and $w_c$ allowed to vary by condition. The trial-wise posterior probability that a response arose from the internal component is:

$$z_i \; = \; P(\text{int} \mid r_i, t_i, c_i) \; = \; \frac{w_{c_i}\, p_{\text{int}}(r_i \mid t_i; a, b)}{w_{c_i}\, p_{\text{int}}(r_i \mid t_i; a, b) + (1 - w_{c_i})\, q_0(r_i)}.$$

**M-step updates**
Mixture weights were updated in closed form:

$$w_c \leftarrow \frac{1}{n_c} \sum_{i:\, c_i = c} z_i,$$

where $n_c$ is the number of stimulus-present trials in condition $c$. The bowtie parameters $(a, b)$ were updated by minimizing the posterior-weighted negative log-likelihood of the internal component:

$$(a, b) \leftarrow \arg\min_{a,b} \left[ -\sum_i z_i \log\, p_{\text{int}}(r_i \mid t_i; a, b) \right], \text{with constraint } a \geq b.$$

Optimization was performed using fminsearch (MATLAB 2020B).

**Initialization and stopping**
Initial $(a, b)$ values were seeded from the observer's best-fitting bowtie parameters estimated from Experiment 1 precision data when available; otherwise we used the sample median across observers. EM iterations were capped at 200 iterations and terminated early when the absolute change in log-likelihood satisfied:

$$\mid \Delta LL \mid < 10^{-6}\, n,$$

(where $n$ is the number of stimulus-present trials used for fitting.)

**Recovered guessing on stimulus-present trials**
To reconstruct guessing structure from stimulus-present trials, we computed a posterior-weighted KDE over stimulus-present responses using the posterior probability of guessing $(1 - z_i)$ as a trial weight:

$$\tilde{q}_{\text{rec}}(r) = \sum_i (1 - z_i) \exp\left[ -\frac{1}{2} \left( \frac{\delta(r, r_i)}{h} \right)^2 \right], \qquad h = 3°,$$

and normalized it to unit mass on the 1° grid:

$$q_{\text{rec}}(r) = \frac{\tilde{q}_{\text{rec}}(r)}{\sum_{r'=1}^{180} \tilde{q}_{\text{rec}}(r')}$$

**Comparison to a uniform-guess null (posterior recovery control)**
Because $q_0(r)$ enters the posterior $z_i$, posterior-weighted recovery could have, in principle, biased $q_{\text{rec}}(r)$ toward $q_0(r)$. To quantify this bias under a null in which stimulus-present guesses are uniform, we simulated stimulus-present responses for

each observer while preserving that observer's fitted internal component and trial/condition structure, but drew guess responses from a distribution $q_{\mathrm{unif}}(r) = 1/180$. We then applied the identical posterior-recovery procedure to obtain $q_{\mathrm{rec,null}}(r)$ and computed its correlation with $q_0(r)$. This simulation was repeated (2000 iterations per observer) to obtain an observer-specific null distribution for the recovered-vs-$q_0$ similarity.

**Recovered guess complementarity with independently measured precision**
Complementarity between stimulus-present recovered guessing and independently measured precision was assessed within each observer as the Pearson correlation between the smoothed recovered guess density $q_{\mathrm{rec,smooth}}$ (from stimulus-present trials) and the smoothed precision landscape $P_{\mathrm{smooth}}$ (from Experiment 1 precision trials), using the same exact circular-shift alignment test described above (left-tailed). Group-level summaries used observer bootstrapping for confidence intervals and a one-sided sign-flip test over observer-level correlations.

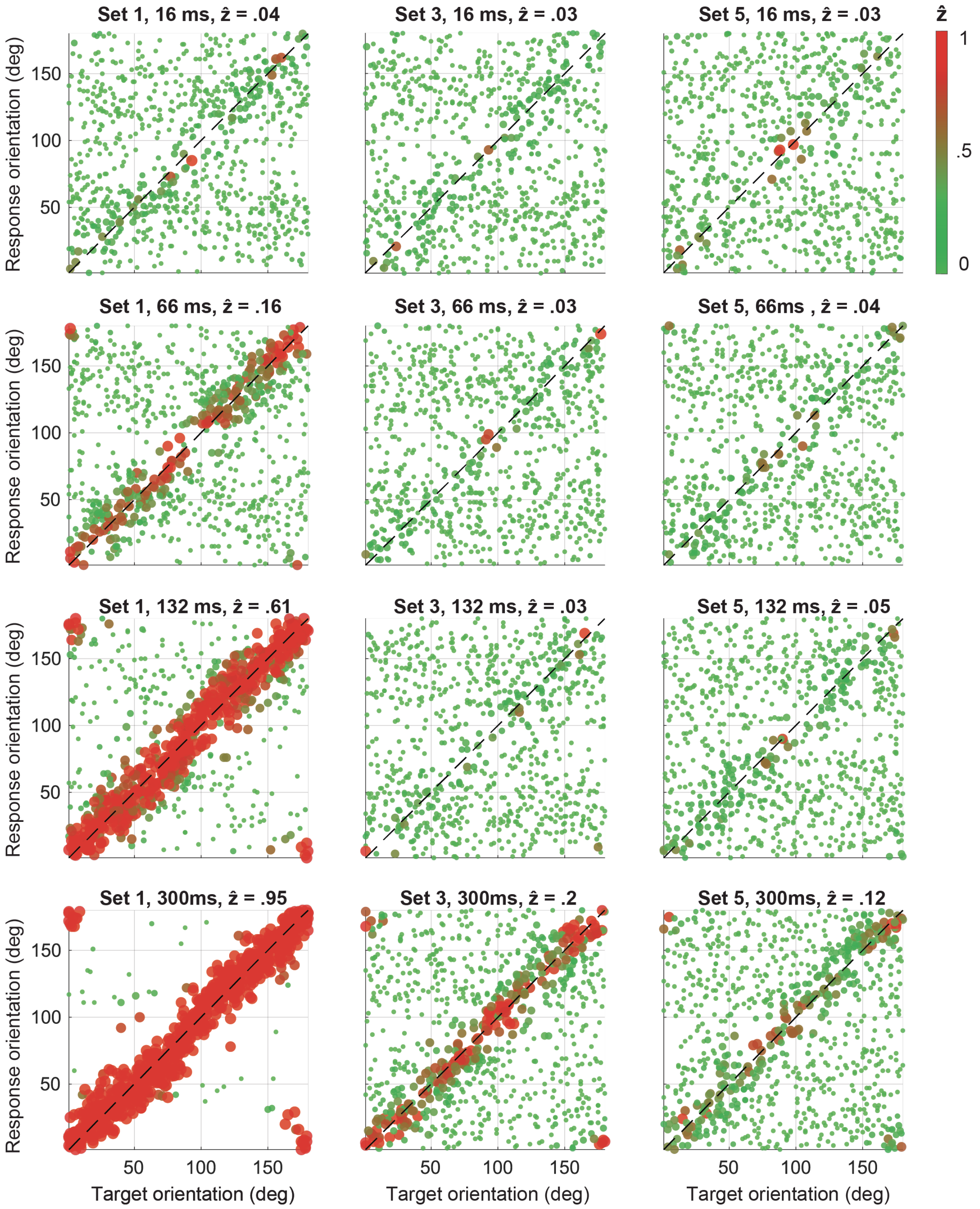


**Supplementary Figure 1. Trial-wise posterior classification ($z_i$) by condition in Experiment 2.** Each panel shows target orientation (x-axis) plotted against response orientation (y-axis) for one set size (columns: 1, 3, 5) and display duration (rows: 16, 66,

132, 300 ms). Points are individual trials, colored by the empirical-guess mixture model's trial-wise posterior probability of the internal (target-driven) component, $z_i$ (color bar). Warmer colors indicate higher $z_i$ (responses more likely generated from the target-centered internal component); cooler colors indicate lower $z_i$ (responses more likely generated from the empirical guess component derived from 0-ms trials). The dashed diagonal indicates perfect target–response correspondence. Panel titles report the mean posterior *z* for that condition.
